# Cecropins contribute to *Drosophila* host defence against fungal and Gram-negative bacterial infection

**DOI:** 10.1101/2021.05.06.442783

**Authors:** A. Carboni, M.A. Hanson, S.A. Lindsay, S.A. Wasserman, B. Lemaitre

## Abstract

Cecropins are small helical secreted peptides with antimicrobial activity that are widely distributed among insects. Genes encoding Cecropins are strongly induced upon infection, pointing to their role in host-defence. In *Drosophila*, four *Cecropin* genes clustered in the genome (*CecA1, CecA2, CecB* and *CecC*) are expressed upon infection downstream of the Toll and Imd pathways. In this study, we generated a short deletion *ΔCec^A-C^* removing the whole *Cecropin* locus. Using the *ΔCec^A-C^* deficiency alone or in combination with other antimicrobial peptide (AMP) mutations, we addressed the function of Cecropins in the systemic immune response. *ΔCec^A-C^* flies were viable and resisted challenge with various microbes as wild-type. However, removing *ΔCec^A-C^* in flies already lacking ten other AMP genes revealed a role for Cecropins in defence against Gram-negative bacteria and fungi. Measurements of pathogen loads confirm that Cecropins contribute to the control of certain Gram-negative bacteria, notably *Enterobacter cloacae* and *Providencia heimbachae*. Collectively, our work provides the first genetic demonstration of a role for Cecropins in insect host defence, and confirms their *in vivo* activity primarily against Gram-negative bacteria and fungi. Generation of a fly line (*ΔAMP14*) that lacks fourteen immune inducible AMPs provides a powerful tool to address the function of these immune effectors in host-pathogen interactions and beyond.

## INTRODUCTION

In the late nineteen seventies, as immunologists were characterizing the antibody immune response of mammals, pioneering studies revealed that insects could resist infection by fearsome human pathogens despite lacking an adaptive immune system. Eventually a landmark discovery by Hans Boman and colleagues showed that insects produced antimicrobial peptides (AMPs) following infection (Steiner *et al*. 1981), invigorating interest in innate immunity (Ganz *et al*. 1985; Lemaitre 2004). These AMPs differed from other previously identified immune effectors in their small size, cationic charge, and amphipathic structure, which allowed them to directly disrupt the negatively charged membrane of microbes. In contrast to another class of well-known immune effectors, the lysozymes, AMPs lack enzymatic activity and require concentrations into the micromolar range to achieve their microbicidal effects (Imler and Bulet 2005; Seo *et al*. 2012; Hanson and Lemaitre 2020). Research has now shown that AMPs are common across the tree of life, with similar molecules contributing to host defence in both plants and animals (Broekaert *et al*. 1995). While they contribute to local defence in barrier epithelia of vertebrates, insect AMPs are most famous for being secreted upon systemic infection from the fat body into the hemolymph, where they reach potent concentrations (Bulet *et al*. 1999). The characterization of a plethora of AMPs with diverse modes of action has enriched our understanding of these immune effectors. However, the functional study of AMPs was limited until recently due to technical challenges in mutating the small AMP genes using traditional genetic approaches. This challenge has now been overcome with the advent of CRISPR/Cas9 technology.

Cecropins were the first inducible AMPs to be isolated, found in the hemolymph of infected pupae of the moth *Hyalophora cecropia* (Lepidoptera) (Hultmark *et al*. 1980; Steiner *et al*. 1981). The helical conformation of Cecropins is thought to promote their interaction with negatively charged bacterial membranes, contributing to pore formation and membrane destabilization, and resulting in the bacterial lysis (Steiner *et al*. 1988). *In vitro* studies have shown that Cecropins have high efficacy against a large panel of gram-negative bacteria at concentrations below the levels induced in insects upon infection (10μM), as well as against some filamentous fungi (DeLucca *et al*. 1997; Ekengren and Hultmark 1999; Ouyang *et al*. 2015). Heterologous expression of *Cecropin* in transgenic rice has also been shown to confer resistance to the rice blast fungus *Magnaporthe oryzae* (Coca *et al*. 2004). Cecropins *in vitro* do not display antimicrobial activities against Gram-positive bacteria (Smeianov *et al*. 2000). However some studies have reported an activity of Cecropins against tumor cells, bacterial biofilms and viruses (Chiou *et al*. 2002; Suttmann *et al*. 2008; Deslouches and Di 2017; Kalsy *et al*. 2020).

AMP regulation and function has been extensively studied in the model insect *Drosophila melanogaster*. The *Drosophila* genome encodes four *Cecropin* genes (*CecA1* and *A2, CecB* and *CecC*) and two pseudogenes (*Cec-Ψ1* and *Cec2*) that are clustered at position 99E2 at the tip of the right arm of the third chromosome (Kylsten *et al*. 1990; Samakovlis *et al*. 1990; Sackton *et al*. 2007). The *Cecropin* locus is adjacent to another gene named *Andropin*, which encodes a related antibacterial peptide expressed in the ejaculatory duct (Samakovlis *et al*. 1991). CecA1 and CecA2 are identical at the protein level, differing only by a few silent mutations at the nucleotide level, suggesting that they emerged from a recent duplication. The four *Drosophila Cecropin* genes are strongly induced in the fat body and hemocytes upon systemic infection. During systemic infection, *Cecropin* genes are regulated mostly by the Imd pathway while also receiving minor input from the Toll pathway (Lemaitre *et al*. 1996; Hedengren *et al*. 1999). Functional studies analyzing the role of Cecropins *in vivo* are scarce. Overexpression of *CecA* in an otherwise *Imd, Toll* immune-deficient background failed to detect a clear protective effect for *CecA* against a battery of pathogens (Tzou *et al*. 2002). Other studies using overexpression approaches have pointed to a role for *CecA* in the regulation of the gut microbiota (Ryu *et al*. 2008). Transgenic mosquitoes overexpressing both Cecropin and Defensin under the control of the vitellogenin promoter displayed an increased resistance to *Pseudomonas aeruginosa* infection, indicating that these AMPs could act cooperatively against this pathogenic bacterium (Kokoza *et al*. 2010).

We have previously generated fly strains with deletions in 10 *Drosophila* AMP genes including: *Defensin*, two *Diptericins (DptA* and *B*), *Drosocin*, four *Attacins (AttA, B, C*, and *D*), *Metchnikowin*, and *Drosomycin* (Hanson *et al*. 2019a). This study revealed that AMPs play an important role in defence against Gram-negative bacteria and also somewhat in defence against fungi. In contrast, another family of host defence peptides with no overt antimicrobial activity *in vitro*, the Bomanins, plays a major role in the elimination of Gram-positive bacteria and fungi (Clemmons *et al*. 2015; Lindsay *et al*. 2018). Importantly, Hanson et al. (Hanson *et al*. 2019a) revealed evidence for synergy and additivity, but also remarkable specificity in the action of AMPs against certain pathogens. However, this study did not address the function of the four Cecropins due to a failure to generate a proper *Cecropin* locus deletion. In the present study, we have generated fly lines carrying a small deletion that removes the four immune *Cecropin* genes, and by using flies carrying this deletion alone or in combination with other AMP mutations, we address for the first time the role of Cecropins in the systemic immune response.

## MATERIALS AND METHODS

### Fly stocks and genetics

The *w^1118^* DrosDel isogenic (iso *w^1118^*) wild type was used as the genetic background for mutant isogenization, as described by Ferreira and colleagues (Ferreira *et al*. 2014). The *ΔCec^A-C^* mutation was generated using CRISPR with two gRNAs and a homology directed repair vector by cloning 5’ and 3’ region-homologous arms into the pHD-DsRed vector, and consequently *ΔCec^A-C^* flies express DsRed in their eyes, ocelli and abdomen. The *ΔCec^A-C^* mutation was generated by Cas9 mediated injection into the iso *Mtk^R1^; Drs^R1^* background. Following this, two rounds of backcrossing were performed to replace the 1^st^ and 2^nd^ chromosome with the *iso DrosDel* 1^st^ and 2^nd^ chromosome, and to recombine the *ΔCec^A-C^* mutation away from other mutations. The resulting stock is here called iso *ΔCec^A-C^*. Afterwards, the *ΔCec^A-C^* mutation was recombined independently with *Drs^R1^* and *AttD^SK1^* on chromosome 3, and introgressed alongside the other AMP mutations on chromosome 2 (Hanson *et al*. 2019a) to generate *ΔAMP14* flies lacking the 14 classical AMP genes from the *Defensin, Drosocin, Attacin, Diptericin, Metchnikowin, Drosomycin*, and *Cecropin* gene families. The iso *Bom^Δ55C^*and iso *Relish^E20^* flies are the same as used in Hanson et al. (Hanson *et al*. 2019a). Finally, the *ΔAMP10* genotype is described in Hanson *et al*.; however we removed the aberrant *Cecropin* locus (*Cec^SK6^*) detected in this line to avoid any potential effects this locus could have on Cecropin-mediated resistance to infection (see Hanson et al. correction notice (Hanson *et al*. 2019b)).

### Microbial culture conditions

Bacteria were grown overnight on a shaking plate at 200 RPM in their respective growth media and at their optimal temperature conditions. They were then pelleted by centrifugation (4000 RPM) at 4°C. The bacterial pellets were diluted to the desired optical density at 600nm (OD_600_).

*Pectobacterium carotovorum carotovorum 15 (Ecc15*) and *Micrococcus luteus* were grown in LB media at 29°C. *Escherichia coli* strain 1106, *Providencia burhodogranariea, Providencia rettgeri* and *Providencia heimbachae* were grown in LB media at 37°C. *Enterococcus faecalis, Listeria monocytogenes* and *Enterobacter cloacae* were cultured in BHI media at 37°C. *Streptococcus pneumoniae* was grown as described by Krejčová and colleagues (Krejčová *et al*. 2019). *Candida albicans* strain ATCC 2001 was cultured in YPG media at 37°C. *Aspergillus fumigatus* was grown at 37°C on Malt Agar; spores were collected in sterile PBS, pelleted by centrifugation and resuspended at the desired OD. *Beauveria bassiana* strain R444 and *Metharizium rileyi* strain PHP1705 commercial spores were produced by Andermatt Biocontrol (products: BB-PROTEC and NOMU-PROTEC).

### Infection experiments and survival

Systemic infections with *P. carotovorum carotovorum 15 (Ecc15*) (Basset *et al*. 2000), *M. luteus, E. coli* strain 1106, *P. burhodogranariea, P. rettgeri, P. heimbachae* (Galac and Lazzaro 2011), *E. faecalis, L. monocytogenes, E. cloacae* and *C. albicans* were performed as followed: 3-5 day old adult females were pricked in the thorax with a 100μm thick needle dipped into a concentrated pellet of bacteria at the desired OD_600_. Infected flies were then maintained at 25 or 29°C and survivals were tracked. Systemic infection with *S. pneumoniae* (Krejčová *et al*. 2019) or *M. rileyi* was performed by injecting 50nL of a concentrated pellet of bacteria or suspension of fungal spores using a nanoinjector and glass capillary needles.

Natural infections with *B. bassiana* were performed by shaking anesthetized flies in a vial with 200mg of spores. Flies were flipped into fresh vials one day after fungal inoculation. A minimum of two replicate survival experiments were performed for each infection with 20 flies per vial on standard fly medium without yeast. Survival was scored daily.

### Bacterial load of flies

Flies were infected (systemic infection) with bacteria at the desired OD_600_. At the indicated time post-infection, flies were anaesthetized using CO_2_, surface sterilized by washing briefly in 70% EtOH, and blotted dry. Pools of 5 flies were transferred in 200μL of sterile PBS and macerated using a pestle. The homogenates were centrifuged at 8,000 RPM for 3 minutes. The supernatants were serially diluted and 7μL droplets (6 replicates) were inoculated on LB agar overnight at 29°C. Colony-forming units (CFUs) were manually counted the following day.

### Gene expression levels by RT-qPCR

Flies that either were unchallenged or were infected systemically by pricking in the thorax with a needle dipped in a pellet of *Ecc15* or *M. luteus* (OD_600_=200) were frozen at −20°C 6h or 12h post-infection, respectively. Gene expression measurements were then performed by RT-qPCR as previously described (Hanson *et al*. 2019a). Briefly, 5 whole flies were homogenized and their RNA was extracted using TRIzol reagent and resuspended in RNase-free water. Reverse transcription was carried out using PrimeScript RT (TAKARA) with random hexamers and oligo dTs. Quantitative PCRs were performed on a LightCycler 480 (Roche) using PowerUp SYBR Green Master Mix.

### Cecropin A injection

Commercially available *Hyalophora cecropia* Cecropin A (Sigma-Aldrich) was diluted in PBS to a concentration of 50μM. Fifty nL of Cecropin was injected into the thorax using a nanoinjector and glass capillary. Flies were left to recover for 2 hours and then pricked with the desired pathogen.

#### MALDI-TOF

Hemolymph samples were collected from either unchallenged flies, or flies pricked with a 1:1 cocktail of *E. coli* and *M. luteus* (OD=200) in 0.1% TFA, as described previously (Hanson *et al*. 2019a). Samples were then added to an acetonitrile universal matrix and subjected to desorption and ionization. Representative spectra are shown. Immune induced peaks were identified based on previous studies (Uttenweiler-Joseph *et al*. 1998; Levy *et al*. 2004) to confirm the absence of AMP-associated peaks, and presence of immune-induced peptides not affected by the various AMP mutations. Spectra were visualized using mMass and figures were additionally prepared using Inkscape v0.92.

### Statistical analysis

Survival analyses were performed using a Cox proportional hazards (CoxPH) multiple comparison model, with Benjamini-Hochberg corrections for p-values, in R 3.6.3. Survival curves included at least one survival experiment with 2 cohorts of 20 flies per treatment. Statistics were represented using a Compact Letter Display (CLD) graphical technique: groups were assigned the same letter if they were not significantly different (p>.05). Quantitative PCR data were compared by one-way ANOVA with Holm-Šídák multiple test correction in Prism R7. Bacterial load values were transformed as log_10_(value+1) to allow graphical representation of the absence (0) of CFUs. Bacterial load data were compared by one-way ANOVA with Holm- Šídák multiple test correction in Prism 7. Statistics were represented using a CLD graphical technique.

## RESULTS

### Generation and characterization of *Cecropin* mutants

We generated a fly line lacking the four *Cecropin* genes, *CecA1, CecA2, CecB*, and *CecC*, which are clustered at 99E and are inducible during the systemic response. For this, we used the CRISPR/Cas9 editing method to generate a 6kb deletion (referred as *ΔCec^A-C^*) that removes the four inducible *Cecropins* but leaves the neighboring *Andropin* gene intact (Fig. 1A). The *ΔCec^A-C^*mutation was generated by Cas9 mediated injection in the *w, Drosdel* (referred to as *w^1118^*) background. The background of the *ΔCec^A-C^* mutation was then cleaned by 2 successive crosses to the *w^1118^ iso* background to remove potential off-target alterations. To confirm the absence of *Cecropin* genes in *ΔCec^A-C^* flies, we performed RT-qPCR of *CecA*1 and *CecC* genes, which are located at the two extremities of the *Cecropin* locus. Expression of *CecA1* or *CecC*, readily observed in the wild type, was not detected in *ΔCec^A-C^* flies (Fig. 1B-C).

**Figure 1:**
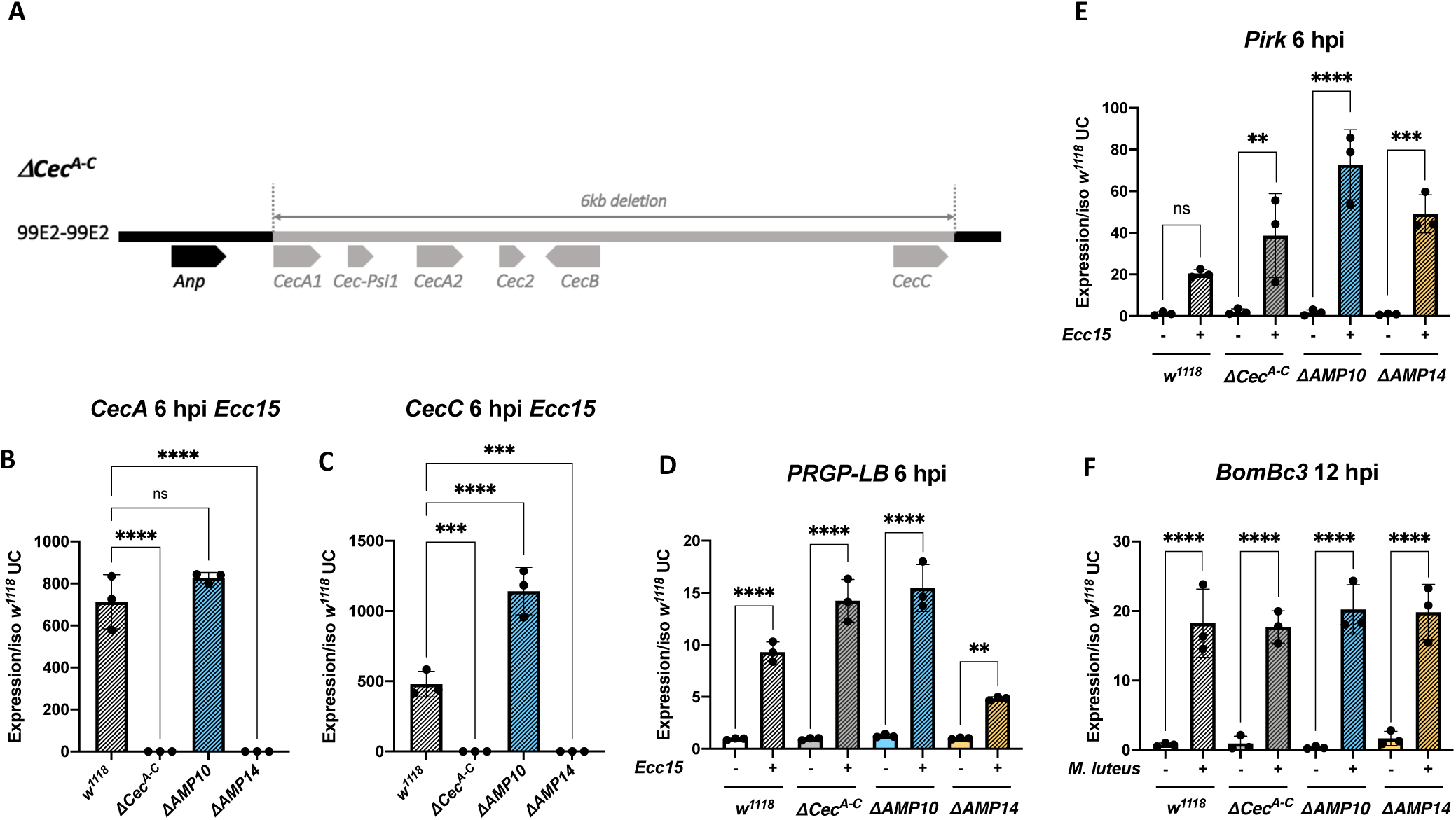
Description and validation of *ΔCecA-C, ΔAMP10* and *ΔAMP14* mutants. (**A**) Schema of the *Cecropin* locus chromosomal deletion removing *CecA1* and *A2, CecB* and *CecC*, plus 2 pseudo genes, *Cec2* and *Cec- Ψ 1* clustered at position 99E2 (III) (**B,C**) RT-qPCR of *CecA* (B) and *CecC* (C) expression in *w^1118^, ΔCec^A-C^, ΔAMP10* and *ΔAMP14* flies 6 hours post *Ecc15* infection. (**D-F**) The Imd (D-E) and Toll (F) pathways are functional in *ΔCec^A-C^, ΔAMP10* and *ΔAMP14* flies after challenge as revealed by expression of target genes upon septic injury with *Ecc15* or *M. luteus. PGRP-LB* and *Pirk* were used as readouts for Imd pathway and *Bomanin (BomBc3*) for Toll pathway. Expression was normalized with *w^1118^* UC to a value of 1. Statistical significance levels were represented as: * = P≤0.05; ** = P≤0.01; *** = P≤0.001; **** = P≤0.0001.

We previously generated a fly line in the *w^1118^ iso* background here referred to as “*ΔAMP10*”, harbouring six mutations that remove ten antimicrobial peptide genes: *Defensin, Metchnikowin*, the four Attacins (*A/B/C/D), Drosomycin*, two Diptericins (*A/B*), and *Drosocin* (Hanson et al., 2019). We recombined the iso *ΔCec^A-C^* mutation with the iso *ΔAMP10* mutations to generate an iso fly line lacking all 14 ‘classical’ antimicrobial peptides (referred to as “*ΔAMP14*”). MALDI-TOF and RT-qPCR analysis confirms the absence of these 14 antimicrobial peptides in *ΔAMP10* and *ΔAMP14* flies (Fig. 1B-C and Fig. S1). The *ΔCec^A-C^, ΔAMP10* and *ΔAMP14* flies were healthy and showed no morphological defects. We also confirmed that the two central NF-κB signaling pathways, Toll and Imd, were functional, as demonstrated by measuring expression of genes characteristic of each of these pathways (Fig. 1D-F). Furthermore, MALDI-TOF proteomic analysis of hemolymph from infected flies 24 hours post infection (hpi) reveals a wild type-like induction of peaks associated with other NF-κB effectors (e.g., Bomanins, Daishos, and Baramicin A) (Fig. S1). Collectively, our study indicates that we have generated a fly line lacking all the Drosophila ‘classical’ AMPs, and that deleting these AMPs does not impact the production of other NF-κB effectors.

### Cecropins contribute to survival against certain Gram-negative bacterial infections

We used wild-type, *ΔCec^A-C^, ΔAMP10* and *ΔAMP14* flies to explore the role that Cecropins play in defence against pathogens during systemic infection. By performing survival analyses with wild-type and *ΔCec^A-C^* flies, we assessed if the absence of the four Cecropins is sufficient to cause an immune deficiency. Likewise, any difference in survival rates between *ΔAMP10* and *ΔAMP14* flies would suggest a contribution of Cecropins that is only apparent in the absence of other AMPs. We first focused our attention on Gram-negative bacterial infections, as cecropins were initially identified for their activity against this class of bacteria, and the four *Cecropin* genes are mostly regulated by the Imd pathway that responds to Gram-negative bacterial infection (Lemaitre and Hoffmann 2007). We challenged wild-type, *ΔCec^A-C^, ΔAMP10* and *ΔAMP14* flies with six different Gram-negative bacterial species, using inoculation doses (given as OD_600_) selected such that Imd deficient, iso *Rel^E20^* mutant control flies were killed. Our survival experiments did not reveal an overt contribution of Cecropins to resistance against the Gram-negative bacteria *Escherichia coli, Pectobacterium carotovorum carotovorum (Ecc15), Providencia rettgeri* or *Providencia burhodogranariea* (Fig 2A-D). In all cases, *ΔCec^A-C^* flies survived as well as wild-type flies, while *ΔAMP10* flies were as susceptible as *ΔAMP14*. One exception was found for *P. burhodogranariea* infection: death of *ΔAMP10* flies was delayed by one day compared to *ΔAMP14* flies, suggesting a contribution of Cecropins in combatting this bacterium early in infection.

**Figure 2:**
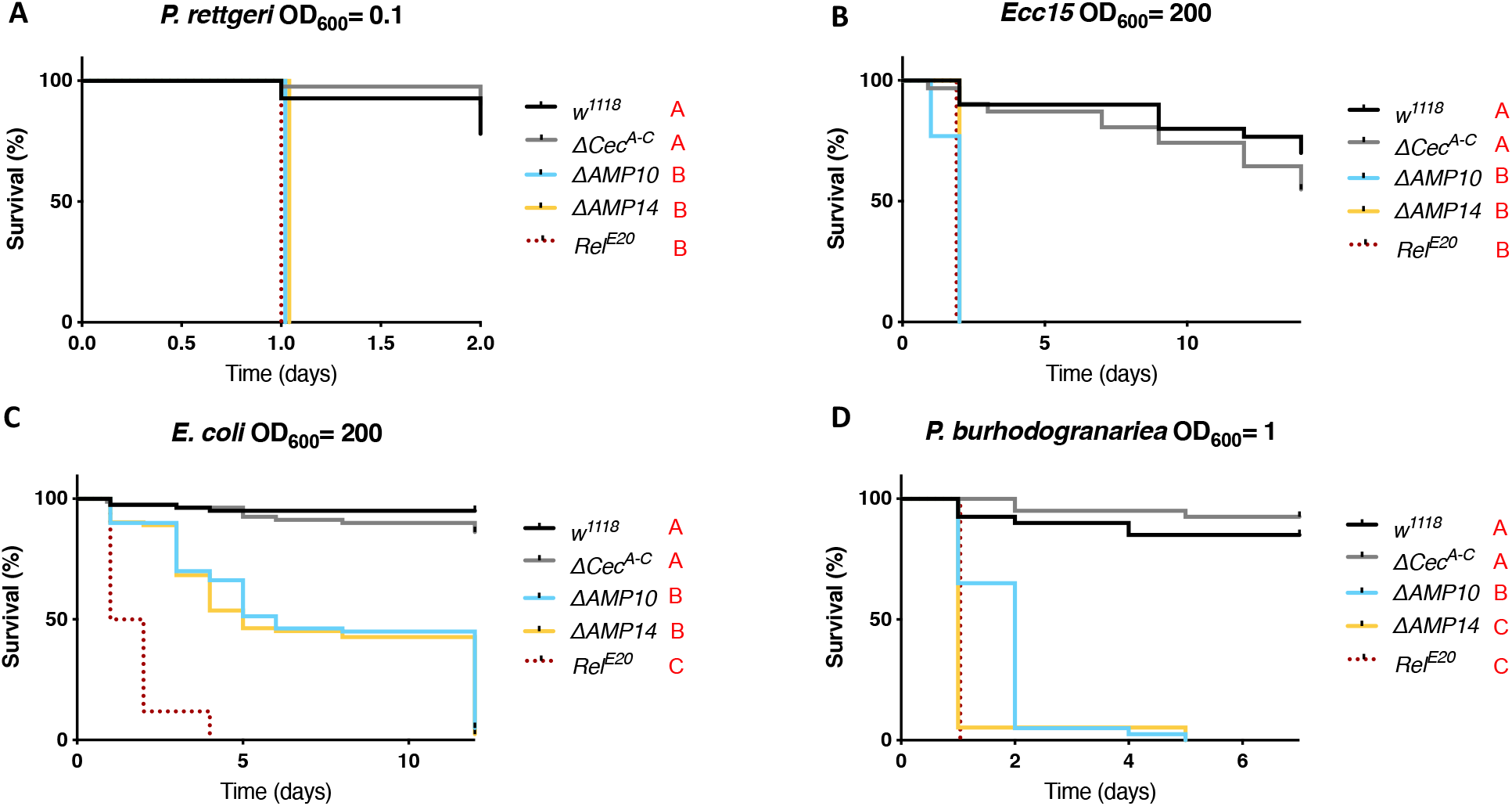
Cecropins are not involved in resistance to a broad spectrum of Gram-negative bacteria. *w^1118^* were used as wild-type flies and *Rel^E20^* as susceptible flies lacking the Imd pathway for all survival experiments to Gram-negative bacterial infection. Female *w^1118^, ΔCec^A-C^, ΔAMP10, ΔAMP14* and *Rel^E20^* flies were pricked in the thorax with an inoculum of (**A**) *P. rettgeri*, (**B**) *Ecc15*, (**C**) *E. coli* or (**D**) *P. burhodogranariea*. Cecropins were not critically involved in combating infection with any bacteria presented here as *ΔCec^A-C^* survived as well as *w^1118^* flies, while *ΔAMP10* and *ΔAMP14* mutant flies died as fast as *Rel^E20^* mutants. Bacterial concentrations are indicated in the figure.

Interestingly, we did identify a prominent role for Cecropins against two Gram-negative bacterial strains: *Enterobacter cloacae and Providencia heimbachae*. Although *ΔCec^A-C^* flies survived *E. cloacae* infection like wild-type flies and *ΔAMP10* flies had only a slight reduction in survival, *ΔAMP14* flies infected with *E. cloacae* instead behaved like *Rel^E20^* mutants lacking Imd signaling entirely (Fig. 3A). This result suggests that the presence of the four *Cecropin* genes confers a mild protective effect against this bacterium in flies that lack ten other AMP genes. A significant difference in CFUs between *ΔAMP10* and *ΔAMP14* flies at 8h post-infection (hpi) confirmed a role of Cecropins in limiting the growth of *E. cloacae* (Fig. 3B). We also observed a consistently higher bacterial load in *ΔCec^A-C^* flies compared to wild-type controls, though this was not significant (p = .18). Moreover, *ΔAMP14* and *Rel^E20^* fly CFUs were similar, consistent with survival data showing complete mortality of these genotypes within 24 hours. These results indicate that the knock-out of the ‘classical’ AMPs in the *ΔAMP14* line fully explains the susceptibility of Imd pathway mutants to *E. cloacae* infection. Next, we attempted to rescue the susceptibility of *ΔAMP10* and *ΔAMP14* flies using commercially available Cecropin A. We injected 50 nL of 50μM *Hyalophora cecropia* Cecropin A (Sigma-Aldrich) or PBS (control) two hours before challenging flies with *E. cloacae*. Interestingly, when we injected Cecropin A two hours prior to *E. cloacae* infection, *ΔAMP10* flies survived as well as wild-type flies previously injected with PBS (Fig. 3C). This result suggests that priming the fly defence by increasing circulating levels of Cecropin is sufficient to combat *E. cloacae* infection, even in flies lacking a broad range of other AMPs. However we did not succeed in rescuing the susceptibility of *ΔAMP14* flies using the same approach. This suggests the rescue effect we observed using *ΔAMP10* flies relies on the total Cecropin levels, which includes both endogenously produced Cecropin and the supplemental Cecropin A we injected. Collectively, our *in vivo* analysis is consistent with previous *in vitro* studies that showed commercial Cecropin A from *Hyalophora cecropia* has activity against *E. cloacae* (Brey and Ashida 1993).

**Figure 3:**
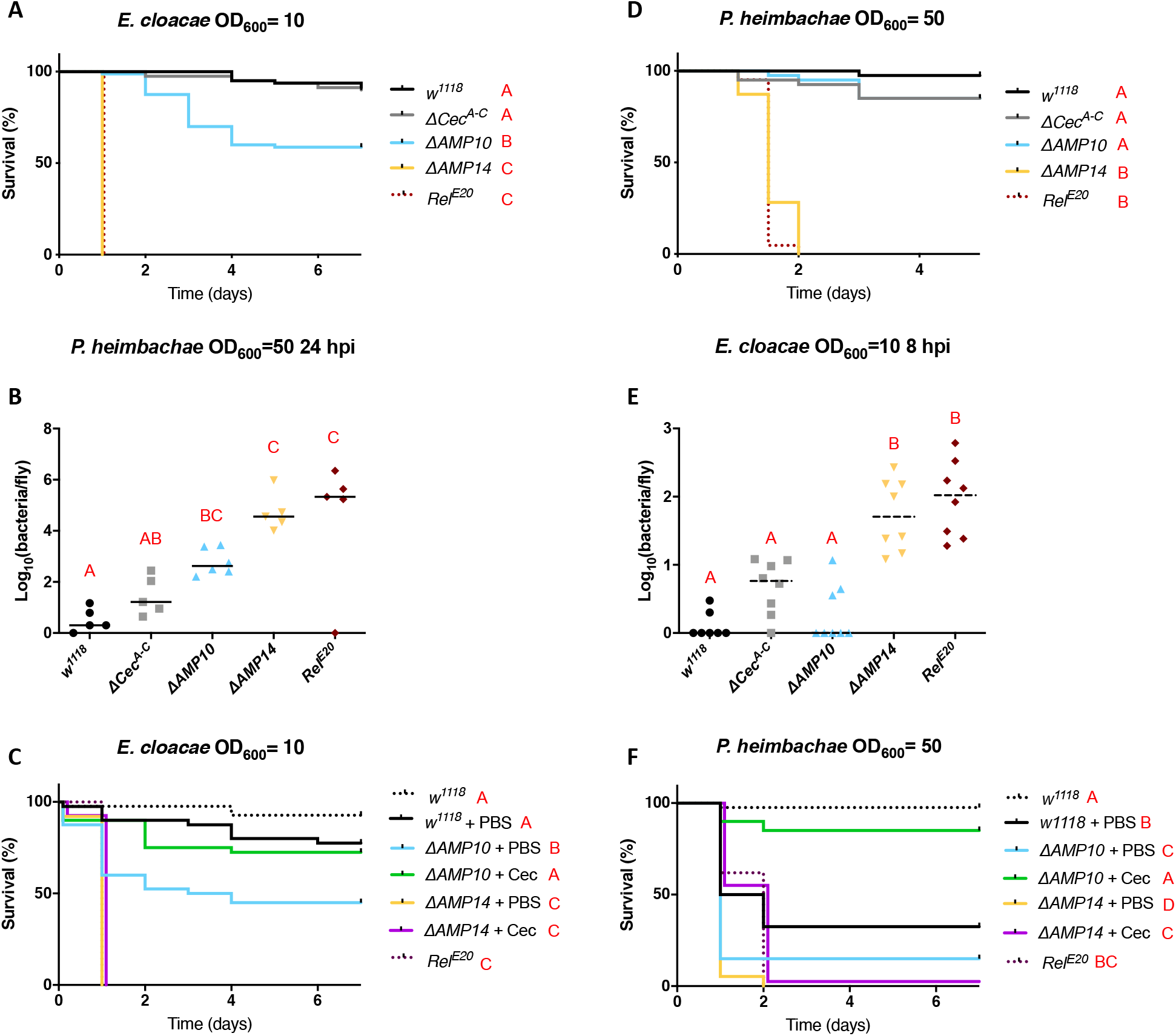
Cecropins are essential in the absence of other AMPs to resist *E. cloacae* and *P. heimbachae* infection. (**A**) Survival experiments upon infection with *E. cloacae* reveal that AMP deficient flies having Cecropins (*ΔAMP10*) are significantly more resistant than those without Cecropins (*ΔAMP14*). (**B**) Bacterial loads (CFU counts) of *w^1118^, ΔCec^A-C^, ΔAMP10, ΔAMP14* and *Rel^E20^* flies 8 hours post infection reveal a significant role for Cecropins in clearing and controlling *E. cloacae*. (**C**) Commercial Cecropin A injection (50 nL at 50 μM) 2 hours prior to *E. cloacae* infection increases the resistance of *ΔAMP10* mutant flies. However, CecA injection did not rescue the susceptibility of *ΔAMP14* flies to *E. cloacae*. (**D-F**) Survival analysis (D) bacterial load measurements 24 hours post infection (E) and Cecropin supplementation experiments (F) in *w^1118^, ΔCec^A-C^, ΔAMP10, ΔAMP14* and *Rel^E20^* flies upon infection with *P. heimbachae* (as described for A-C).

Similarly, we observed a contribution of Cecropins against the Gram-negative bacterium *P. heimbachae* in flies lacking other AMPs (fig. 3D-F). While *ΔAMP10* flies were able to survive this infection as wild type at OD_600_=50, *ΔAMP14* flies again behaved like *Rel^E20^* mutants and suffered complete mortality (Fig. 3D); the *ΔCec^A-C^* mutation alone did not increase susceptibility. Bacterial load measurement performed on flies collected 24 hpi revealed a contribution of Cecropins both in the presence and absence of other AMPs (Fig. 3E). We again injected commercial *H. cecropia* Cecropin A in an attempt to rescue the susceptibility of *ΔAMP10* and *ΔAMP14* flies to *P. heimbachae* (Fig. 3F). Using this bacterial infection model, previous injection of PBS increased the susceptibility of wild-type flies to *P. heimbachae*. Strikingly, however, injection of Cecropin A prior to infection rescued survival of *ΔAMP10* flies to a level comparable to uninjured wild-type flies. Moreover, injection of *H. cecropia* Cecropin A improved the survival of *ΔAMP14* flies against *P. heimbachae*, extending *ΔAMP14* lifespan compared to PBS controls.

In summary, our results reveal that Cecropins contribute to *Drosophila* host defence against a subset of Gram-negative bacteria, and that this contribution is more readily apparent when other AMPs are also lacking.

### Cecropins are not involved in the resistance to Gram-positive bacteria

Previous work with the *ΔAMP10* flies did not reveal a role of *Drosophila* AMPs against Gram-positive bacteria, indicating instead that other immune effectors – notably the bomanins – play a predominant role against this class of microbes (Hanson *et al*. 2019a; Lin *et al*. 2020). Therefore, we were curious if the added loss of Cecropins would reveal a cryptic contribution of *Drosophila* AMPs to defence against Gram-positive bacteria. For this, we challenged wild-type, *ΔCec^A-C^, ΔAMP10* and *ΔAMP14* flies with three Gram-positive bacteria: *E. faecalis, S. pneumoniae* and *L. monocytogenes* (Fig. 4A-C). *E. faecalis* and *S. pneumoniae* contain Lysine-type peptidoglycan that is known to predominantly activate the Toll pathway while *L. monocytogenes* has DAP-type peptidoglycan, and is known to activate both the Toll and Imd pathways (Leulier *et al*. 2003). In these experiments, we included iso *Bom^D55C^* control flies, which lack ten *Bomanin* genes and are known to be susceptible to Gram-positive bacterial and fungal infections (Clemmons *et al*. 2015). Our survival experiments did not reveal a major role of Cecropins individually or alongside other AMPs in combating these Gram-positive bacterial species, but confirmed the importance of bomanins (Fig. 4A-C).

**Figure 4:**
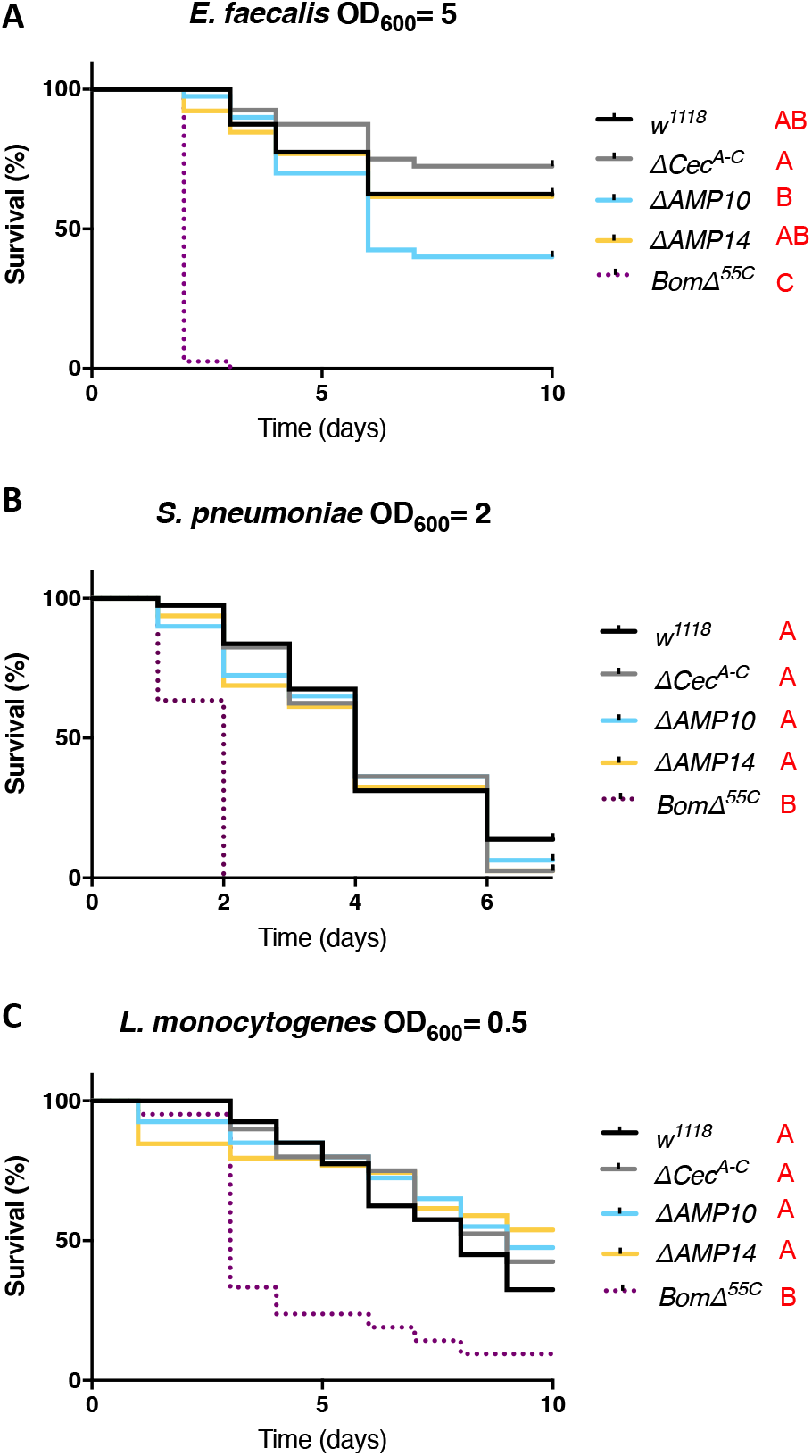
Cecropins are not involved in resistance to Gram-positive bacteria. *w^1118^* were used as wild-type flies and *BomD^55C^* as susceptible flies lacking 10 *Bomanin* genes for all survival experiments to Gram-positive bacterial infection. *w^1118^, ΔCecA-C, ΔAMP10, ΔAMP14* and *BomD^55C^* flies were pricked in the thorax with an inoculum of (**A**) *E. faecalis*, (**B**) *S. pneumoniae* and (**C)** *L. monocytogenes*. Cecropins were not involved in combating infection of 3 bacterial species: *ΔCec^A-C^, ΔAMP10* and *ΔAMP14* flies survived as well as *w^1118^* flies. Bacterial concentrations are indicated in the figure.

### Cecropins can contribute to antifungal defence

While Cecropins were initially identified as antibacterial peptides, further *in vitro* studies have suggested an antifungal activity as well (Ekengren and Hultmark 1999; Andrä *et al*. 2001). We therefore investigated the contribution of Cecropins to resistance upon septic injury with any of four fungal species: the entomopathogenic fungi *Metarhizium rileyi* and *Beauvaria bassiana* R444, the opportunistic mould *Aspergillus fumigatus*, and the yeast *Candida albicans*. Survival analysis did not reveal a prominent role for AMPs or Cecropins against *M. rileyi, A. fumigatus* or *C. albicans*, as *ΔCec^A-C^, ΔAMP10* and *ΔAMP14* flies generally survived as the wild-type (Fig. 5A-C). However, *ΔAMP14* flies suffered greater mortality to *B. bassiana* using a natural infection mode, compared to wild type flies (Fig. 5D). We additionally introduced *B. bassiana* spores directly into the hemolymph by septic injury for more controlled fungal infection kinetics, and measured fungal load at 48h hpi by RT-qPCR. Monitoring pathogen load revealed that in *ΔAMP14* flies, *B. bassiana* loads were higher than levels found in wild type (p=.07) and *ΔAMP10* flies (Fig. 5E). Taken together, these results show a contribution of Cecropins to defence against the entomopathogenic fungus *B. bassiana*.

**Figure 5:**
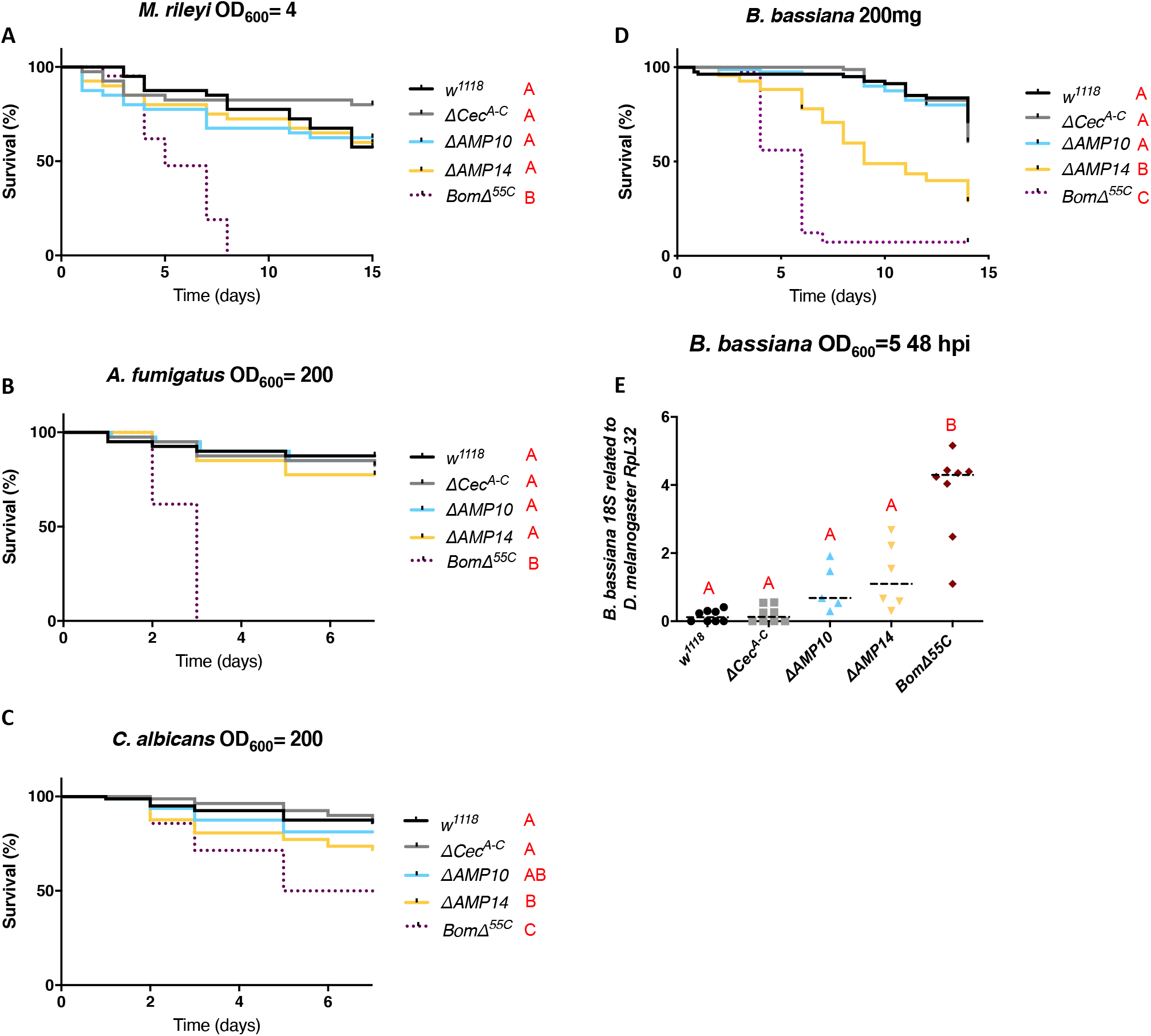
Cecropins contribute to antifungal defence against *B. bassiana*. (**A-C**) *w^1118^* were used as wild-type flies and *Bom^D55C^* as susceptible flies lacking 10 *Bomanin* genes for all survival experiments to fungal infections. Cecropins were not involved in combating infection of (**A**) *M. rileyi*, (**B**) *A. fumigatus* and (**C**) *C. albicans*, as *w^1118^, ΔCec^A-C^, ΔAMP10* and *ΔAMP14* flies survived as well as *w^1118^* flies. (**D**) Survival upon (**D**) natural infection with *B. bassiana* reveals a significant increase in resistance of *ΔAMP10* flies compared to *ΔAMP14* flies, suggesting an important role for cecropins in combating *B. bassiana*. (**E**) *B. bassiana* load (measured by *B. bassiana* 18S expression related to *D. melanogaster RpL32*) is increased (p = .07) in *ΔAMP14* flies compared to *w^1118^, ΔCec^A-C^* and *ΔAMP10* flies 48 hours post septic infection. Fungal concentrations are indicated in the figure.

## DISCUSSION

In this study, we generated flies lacking the four-immune inducible *Cecropin* genes to address their function alone or in combination with other *AMP* gene mutations. *ΔCec^A-C^* and *ΔAMP14* flies were viable, fertile and did not show any morphological defect. Moreover, they display normal activation of the Imd and Toll pathways, suggesting that the classical *Drosophila* AMPs do not contribute to immune signaling, in contrast to mammalian AMPs (Mookherjee *et al*. 2020).

Our survival analyses reveal a role of Cecropins in the defence against certain Gram-negative bacterial species. However, we could not identify a bacterial species or context for which flies mutant for *Cecropin* genes alone succumb faster than wild-type. Studies of other AMPs have revealed that certain AMPs exhibit a high degree of specificity in determining host-pathogen interactions, as illustrated by the requirement of *Diptericin* in defence against *P. rettgeri*, *Drosocin* in defence against *E. cloacae*, and the recently-described *Daisho* and *Baramicin A* genes in defence against *Fusarium oxysporum* and *Beauveria bassiana* fungi, respectively (Unckless *et al*. 2016; Hanson *et al*. 2019a, 2021; Cohen *et al*. 2020). Further studies may reveal bacteria for which the presence of Cecropins is essential for survival.

The most striking phenotype in the present study was that loss of Cecropins has a marked effect on *E. cloacae* and *P. heimbachae* infection in flies also lacking other AMP genes. As such, we reveal an important but cryptic contribution of Cecropins in defence against these bacteria. Generation of flies lacking refined subsets of AMPs might narrow down the specific groups of peptides key to defence against *E. cloacae* and *P. heimbachae*. The enhanced growth of *E. cloacae* in AMP mutants that also lack Cecropins is a particularly striking demonstration of their importance. In this infection model, the presence of Cecropins dictates whether AMP mutant flies initially suppress bacterial growth, or phenocopy *Rel^E20^* flies deficient for Imd signaling. Cecropins are induced with faster kinetics than most other AMPs, with a peak expression as early as 3hpi (Lemaitre *et al*. 1997; De Gregorio *et al*. 2002). As *Cecropins* encode simple helical peptides that do not require extensive post-translational modification, it is tempting to speculate that they become functional more rapidly, and play an important role in combatting bacteria specifically at this early phase of infection.

Our study also reveals that endogenous Cecropins can play a role in defence against fungi, but not against Gram-positive bacteria. Thus, our *in vivo* study corroborates the antifungal and antibacterial activities of Cecropins previously observed with *in vitro* studies (Samakovlis *et al*. 1990; DeLucca *et al*. 1997; Ekengren and Hultmark 1999). While Imd is crucial for the expression of the four Cecropins, the Cecropin response to infection relies in part on Toll signaling (De Gregorio *et al*. 2002). As such, the contribution of Cecropins to defence against fungi could help explain the regulation of the *Cecropin* locus by both Toll and Imd pathways.

The observations that AMP genes are induced to an incredible extent, reach high peptide concentrations in the hemolymph, and display *in vitro* microbicidal activity are all consistent with their role as immune effectors. Use of both *ΔAMP10* and *ΔAMP14* flies has confirmed the important contribution of AMPs to host defence against Gram-negative bacteria and fungi, but not against Gram-positive bacteria. *Drosophila* AMPs also regulate the gut microbiota downstream of the Imd pathway, a function consistent with their bactericidal activity (Marra *et al*. 2021). However, recent studies have suggested that AMP-like genes may play more subtle roles in other processes like memory formation (Barajas-Azpeleta *et al*. 2018), an erect wing response upon infection (Hanson *et al*. 2021), tumor control (Parvy *et al*. 2019; Araki *et al*. 2019), or regulation of JNK signaling in the salivary gland (Krautz *et al*. 2020). While we confirm a primary importance for Cecropins and other AMPs in the systemic immune response, exploring the functions of AMPs including Cecropins in non-canonical roles is an exciting future direction of research.

Our study and others contribute to the rapid progress made towards understanding the roles of *Drosophila* immune effectors. Research on the effector response has stagnated for over a decade, but recent functional characterizations by loss of function of key effectors (*Cecropins, Defensin, Attacins, Diptericins, Drosocin, Drosomycin, Metchnikowin, Bomanins, Daishos*, and *Baramicin*) has greatly advanced our understanding of the roles of these effectors (Lindsay *et al*. 2018; Hanson *et al*. 2019a; Cohen *et al*. 2020; Huang *et al*. 2020). Most importantly, these studies correct the assumptions of the previous “cocktail” model for AMP-pathogen interactions (Yan and Hancock 2001; Lazzaro 2008; Zdybicka-Barabas *et al*. 2012; Rahnamaeian *et al*. 2016), revealing some AMPs to be general effectors against most pathogens, while others act as “silver bullets” specifically required for defence against certain pathogens. The susceptibility of Toll and Imd pathway mutants to specific pathogens can now be directly linked to the susceptibility of mutants for immune effectors regulated by these pathways (Hanson and Lemaitre 2020). As new genetic techniques allow greater characterization of the roles of known immune effectors, many of them yet remain to be characterized, notably a number of short peptide genes highlighted by old and recent transcriptomic studies (Gregorio *et al*. 2001; Troha *et al*. 2018; Tattikota *et al*. 2020; Schlamp *et al*. 2021). However, we are likely still exploring inside the box when assuming a unique immune role for these peptides.

Our study also highlights the power of multiple mutation analysis, as the role of *Cecropins* would not have been uncovered *in vivo* by mutating individual genes. While we have begun exploring the combinatorial potential of AMPs in defence against infection, future studies will benefit from probing the interaction of immune effectors like AMPs with other mechanisms of host defence such as phagocytosis or melanization. With the advent of CRISPR/Cas9 technology and many recently described mutants, the interactions of AMPs in defence are just the tip of the iceberg in developing a global framework to understand the *Drosophila* immune response.

## ACKNOWLEDGEMENTS

We are thankful to EPFL Proteomics Core Facility for help on the MALDI-TOF analysis. Brian Lazzaro generously provided *Providencia* species (Galac and Lazzaro 2011) and Adam Bajgar (Krejčová *et al*. 2019) provided *Streptococcus pneumoniae* species used in this study. We thank Hannah Westlake for editing of the manuscript. We are grateful for the SNF Sinergia grant CRSII5_186397 for the funding of this study.

---

**Figure S1: *ΔAMP14* flies retain the induction of non-AMP Immune Molecules (IMs) upon systemic infection.** Pools of ~40 mixed male and female flies were pricked with a 1:1 mix of OD_600_=200 *E. coli* and *M. luteus* bacteria, and ~40nL of total hemolymph was extracted by nanoject 24hpi, and directly placed in 5uL of 0.1% TFA. Peptide samples were run on an acetonitrile matrix as described previously (Hanson *et al*. 2019a). Hemolymph from infected *w^1118^* wild-type in blue (top) and *ΔAMP14* flies in pink (bottom) are shown. Peaks corresponding to AMP products in the wild-type are labelled in pink, including IM7 whose sequence is unknown, but is absent in *ΔAMP14* flies. Of note, a novel immune-induced peak was observed at ~3027 Da in *AMP* mutants. This peak corresponds to an unknown peptide that is masked by Metchnikowin (~3026 Da) in the wild-type. Reflectron analysis confirms a different spectrum profile for this peak. *Drosophila* IMs are described in (Levy *et al*. 2004) and (Uttenweiler-Joseph *et al*. 1998)

