## Supplementary figures and images for "Cecropins contribute to *Drosophila* host defence against fungal and Gram-negative bacterial infection" (from another version of this preprint: version 1, 2021-05-06)

### Supplemental Figure 1

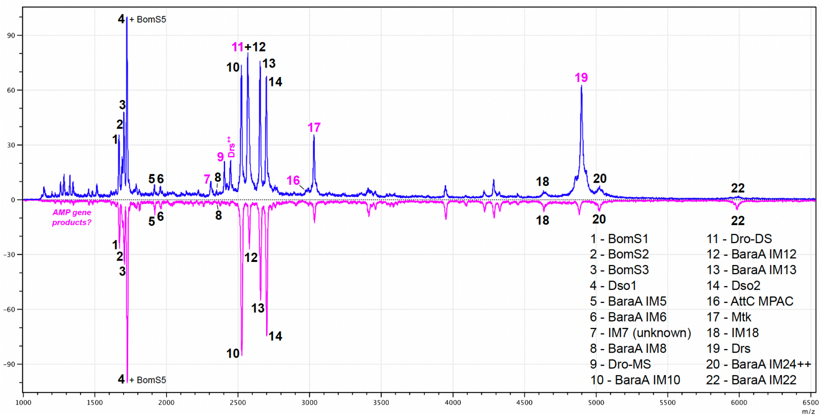
